# Modeling land use impacts on spatial distribution of floating plastic litter in surface water of Southeast Asian archipelago

**DOI:** 10.1101/2021.02.17.431695

**Authors:** Andri Wibowo

## Abstract

Plastics are present in many ecosystems including floating in surface water of remote archipelago and this can lead to the increase in plastic litter density. Whereas the spatial model of plastic litter density related to the population inhabits isolated archipelago is still limited. And what are the underlying factors driving the presence of plastic litter is also poorly understood. This study is trying to find the answers of those questions. The study was implemented in Thousand Island archipelago located in North of Java Island, one of populated islands in Southeast Asia. The studied surface water covers an area of 10000 Ha and consists of 10 islands with 3 islands are occupied by settlements and the remaining islands are occupied by vegetation. This study has recorded 3 types of floating macro-litter from water that consist of PET, HDPE, and LDPE litter. The plastic litter was observed concentrated in the east sides of archipelago where the populated islands were located. The spatial models show LDPE litter was distributed in the vast areas in comparison to PET and HDPE litter. Beside land use variables, the model has confirmed that the population density was the main underlying factors contribute to the plastic litter density in Thousand Island archipelago. The model can be applied to estimate PET (AIC = −0.53060) and HDPE (AIC = 18.28828) litter density. While LDPE litter density was influenced by population (AIC = 22.60201) rather than population density factors.

## INTRODUCTION

Plastic litter is the anthropogenic products and the magnitude of plastic litter mainly in environment is related to the anthropogenic occupation along the natural habitats (van Emmerik & Schwarz 2019). Aydin et al. (2016) have reported that land use types of agricultural and industrial have contributed to 6% plastic litter discharges in the environment. Most plastic litter presence was reported in inland ecosystems and inshore coastal areas rather than from offshore. While recently, the presences of plastic litter in the remote islands have been reported. Plastic has reached from pristine Western Indian Ocean (Dunlop et al. 2020). to the isolated islands in the Pacific. Jennifer et al. (2017) have reported the density of plastic debris in uninhabited island was the highest with density up to 671.6 items/m^2^ (mean ± SD: 239.4 ± 347.3 litter items/m^2^) on the surface of the beaches. Considering the recent vast distribution of plastic litter, this study aims to assess the floating plastic litter density mainly in surface water surrounding the remote archipelago in Southeast Asia regions.

## MATERIALS AND METHODS

### Study area

The study area was an archipelago located in the north of Java Island, Indonesia. This archipelago was named as Thousand Island owing to the numbers of islands in this archipelago that equals to 345 islands whereas only 11 islands that were occupied. Those islands support the livelihood of 24295 people. The study area was focusing on the central areas of the archipelago water within areas of 10000 Ha. In these core areas, there are 10 selected islands where the sampling stations were located. Those islands were not occupied except islands or sampling stations number 6, 7, and 9. The island number 7 is occupied by 2289 people and 1004 people living in island number 9. The island number 7 was smaller than island number 9 and has the population density of 254.33 people/Ha and 62.75/Ha people for island number 9. The land use classifications for those islands were vegetation and settlement for island number 6, 7, and 9. The sampling activities to collect plastic litter from surface water were located in the north, east, south, and west sides of those islands.

### Method

#### Plastic litter surveys

In each sampling station in selected 10 islands, macro-litter plastic (sizing > 5 mm) surveys will be conducted using methods by Carson et al. (2011) and Aytan et al. (2019). Presences of floating plastic litter in surface water will be recorded in sampling quadrats sizing 10 m × 10 m (100 m^2^) and litter was measured as number of plastic litter numbers/100 m^2^. Each collected plastic litter will be identified and classified into its plastic polymer type including PET (plastic bottle), HDPE (plastic wrap), and LDPE (plastic bag). In each sampling station, survey using sampling quadrats sizing 100 m^2^ will be replicated 3 times to obtain representative data.

#### Spatial and land use model

Spatial model used in this paper is addressed to assess the distribution of identified PET, HDPE, and LDPE plastic litter densities in surface water. The plastic litter spatial model was developed using interpolation method. The obtained plastic litter values were tabulated into GIS table and estimated using inverse distance weight method. The plastic litter spatial model was denoted as plastic litter numbers/100 m^2^ and presented as continuous data. The plastic litter data were overlayed with the island land use thematic layers. Those islands selected as sampling stations were classified into vegetation and settlement land uses.

#### Akaike plastic litter model selection

This activity is aiming to assess the anthropogenic factors with their variables that contribute more to the floating plastic litter density in archipelago consists of PET, HDPE, and LDPE densities. Plastic litter density model as function of anthropogenic factors including population and population density was developed using Akaike Information Criterion (AIC). The AIC was developed using the linear regression. The measured parameters included in AIC are R^2^ and adjusted R^2^. To build the model, 2 explanatory covariates including population and population density were included in the analysis.

#### Spatial model of plastic litter

The Figure 2, 3, and 4 presents the spatial distribution of each PET, HDPE, and LDPE litter density floating across the surface water of Thousand Island archipelago covering water sizing 10000 Ha. In surface water, the LDPE and HDPE litter were more common than PET litter. The densities of LDPE and HDPE litter were 5 items/100 m^2^. This density was almost 10 fold larger than PET litter density. Surface water of Thousand Island archipelago only contains 0.5 PET items/100 m^2^. From the model, it is clear that the density patterns of plastic litter are following the land use patterns of those islands. Most plastic litter was concentrated in the east sides of the sampling areas where most inhabited islands were located. These islands included sampling stations number 6, 7, and 9. The west sides of the archipelago were dominated by islands that were inhabited by vegetation as can be observed in sampling station numbers 1, 2, 3, 4, and 5. In these areas, the density of plastic litter was lower. In the east side, the spatial model shows LDPE litter was distributed in the vast and greater areas than PET and HDPE litter. Whereas, there are several uninhabited islands that their surface water were influenced by the presence of the plastic litter as estimated by the spatial model. Those are sampling stations number 8 and 10. The estimate presence of plastic litter in those uninhabited islands is related to their geographical positions that are closed to the inhabited islands.

**Figure 1.**
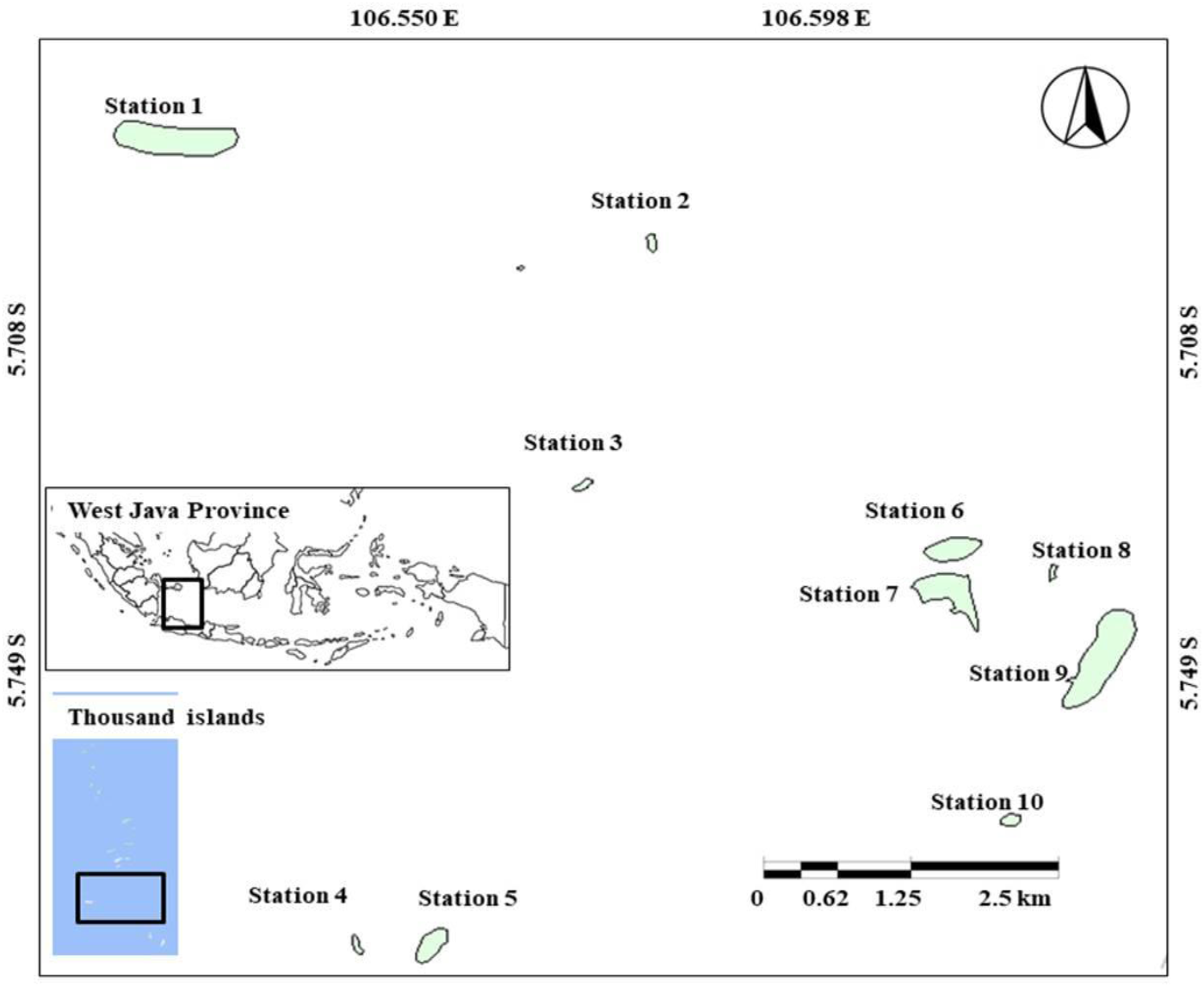
Location of study area and 10 sampling stations in Thousand Island archipelago, Indonesia.

**Figure 2.**
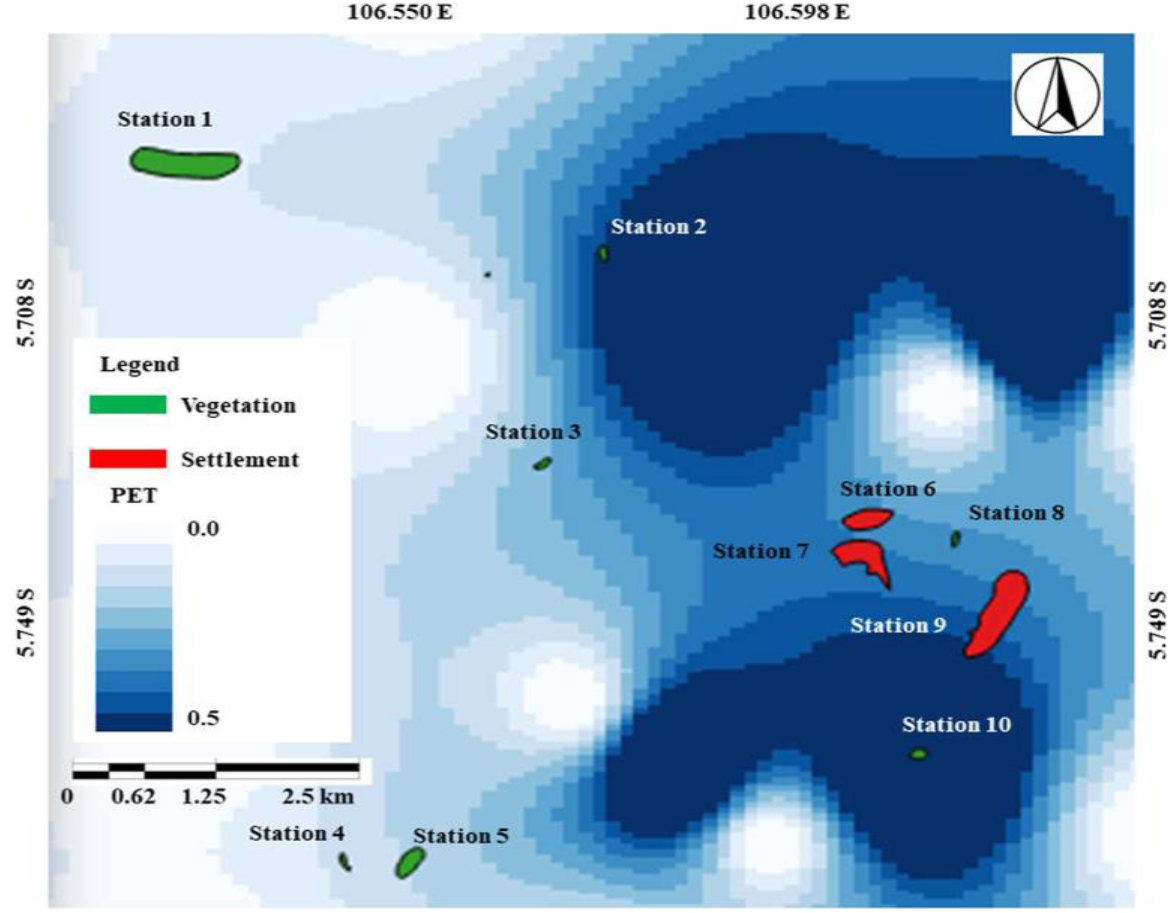
PET litter density (items/100 m^2^) spatial model in surface water of Thousand Island archipelago Indonesia

**Figure 3.**
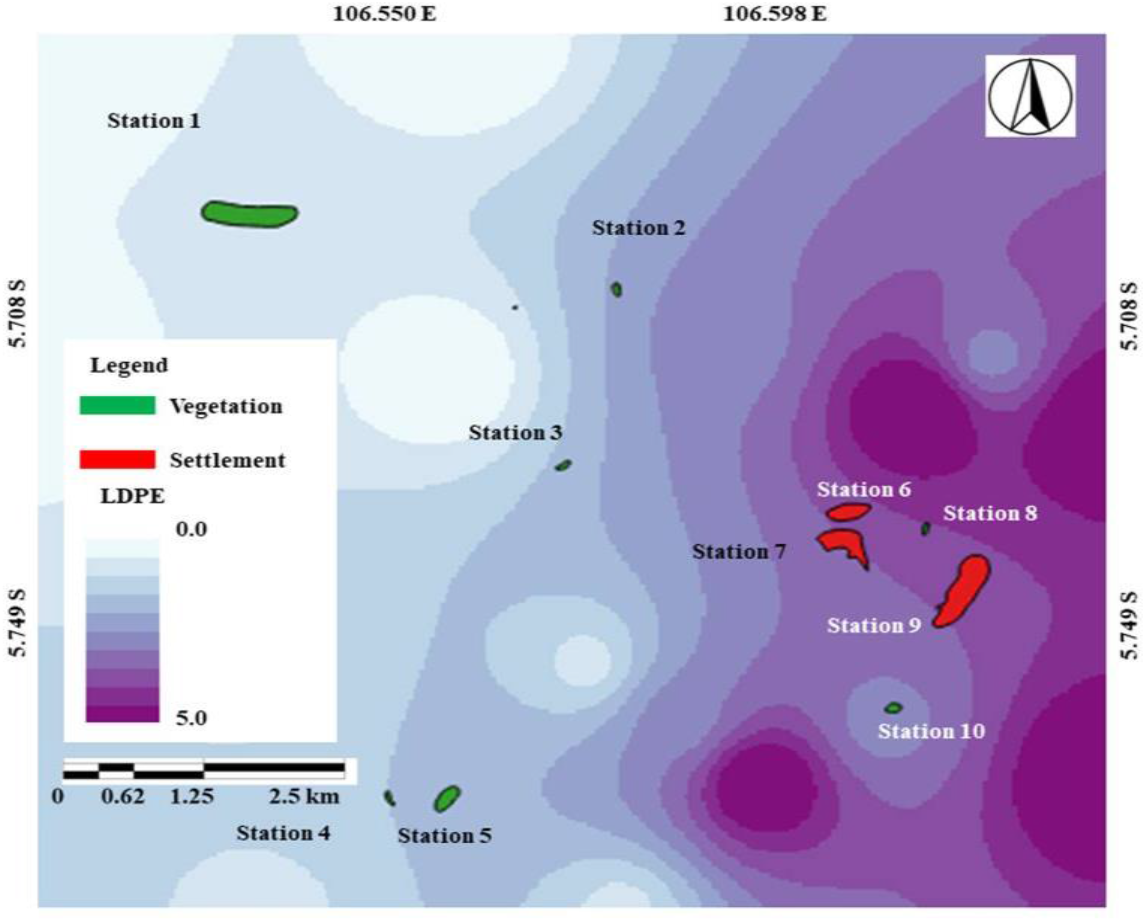
LDPE litter density (items/100 m^2^) spatial model in surface water of Thousand Island archipelago

**Figure 4.**
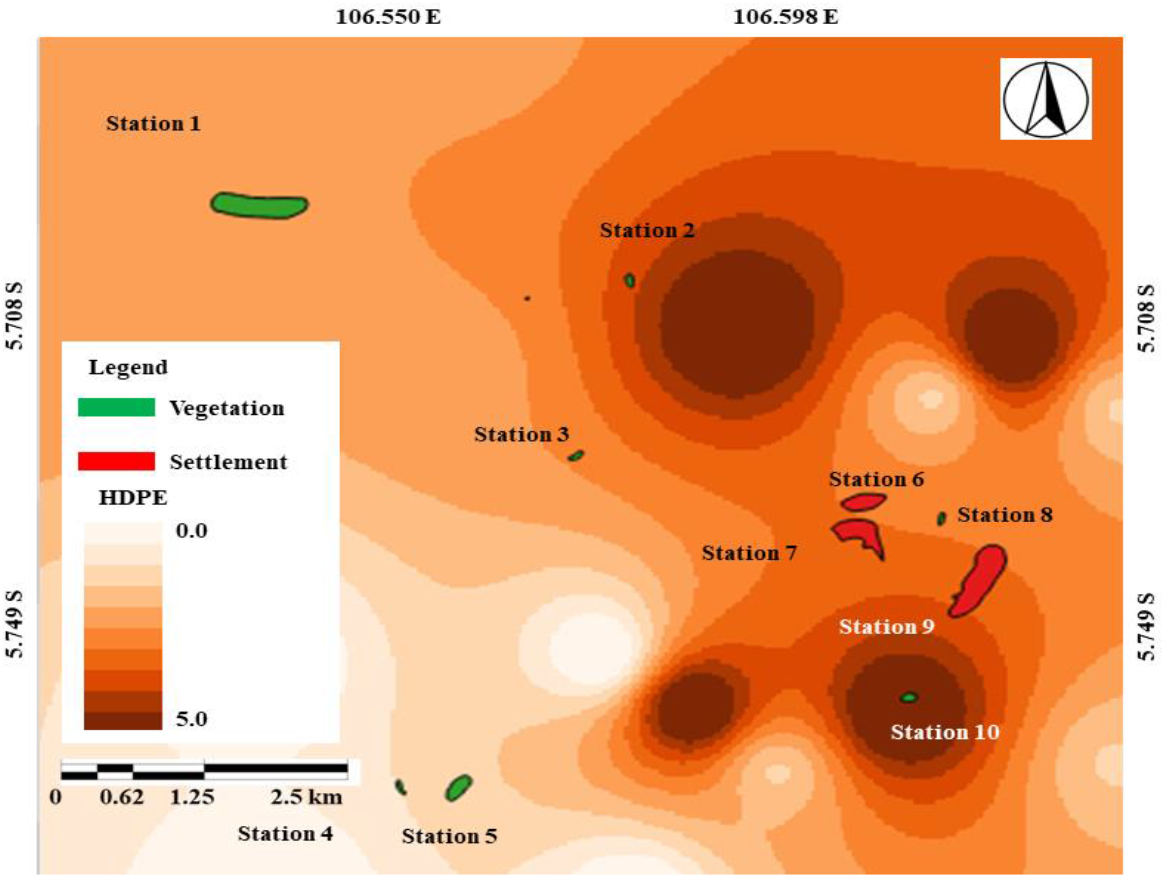
HDPE litter density (items/100 m^2^) spatial model in surface water of Thousand Island archipelago

**Figure 5.**
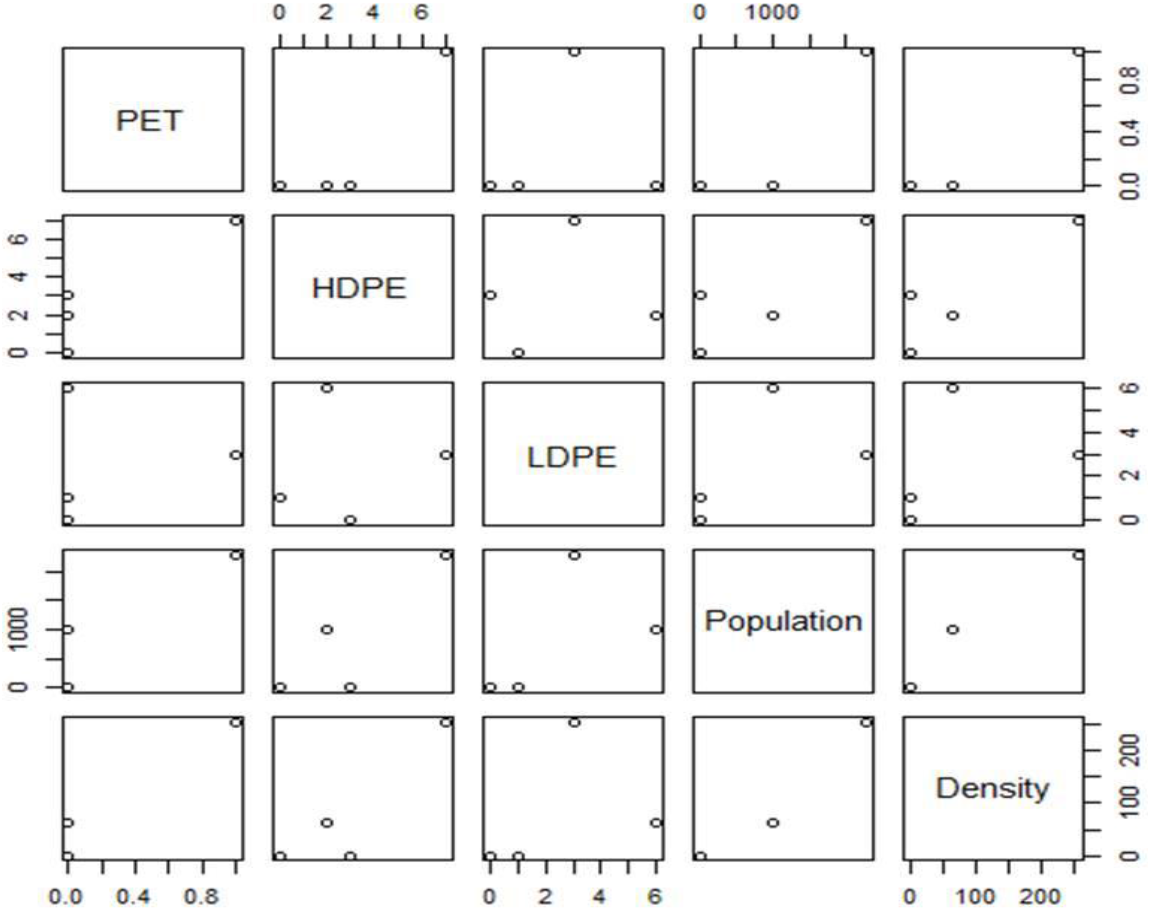
Correlations of each PET, HDPE, and LDPE plastic litter density with population and population density covariates

#### Population and population density

There are 3 islands in the Thousand Island archipelago that are occupied by people. Whereas those islands have differences in areas and numbers of inhabitants. The sampling station number 7 is the island that has the highest population density. In comparison to other inhabited islands, this island has the smallest areas while the population is the highest.

#### Plastic litter model selection

Akaike model selection confirms that the population density variable contributes more to the density of plastic litter mainly PET and HDPE in surface water. This is indicated by the lower AIC values (Table 1). While as can be seen from PCA in Figure 6, the LDPE litter density is influenced more by population rather than density variables. This is also confirmed by the lower AIC of 22.60201.

**Table 1.** Akaike model selections for each PET, HDPE, and LDPE plastic litter density with population and population density covariates

| Model | AIC | R<sup>2</sup> | Adj R<sup>2</sup> |
| --- | --- | --- | --- |
| PET~(population) | 4.013044 | 0.81 | 0.715 |
| PET~(density) | -0.53060 | 0.939 | 0.9085 |
| HDPE~(population) | 19.70776 | 0.7227 | 0.5841 |
| HDPE~(density) | 18.28828 | 0.8056 | 0.7083 |
| LDPE~(population) | 22.60201 | 0.2922 | -0.06169 |
| LDPE~(density) | 23.41536 | 0.1326 | -0.3011 |

**Figure 6.**
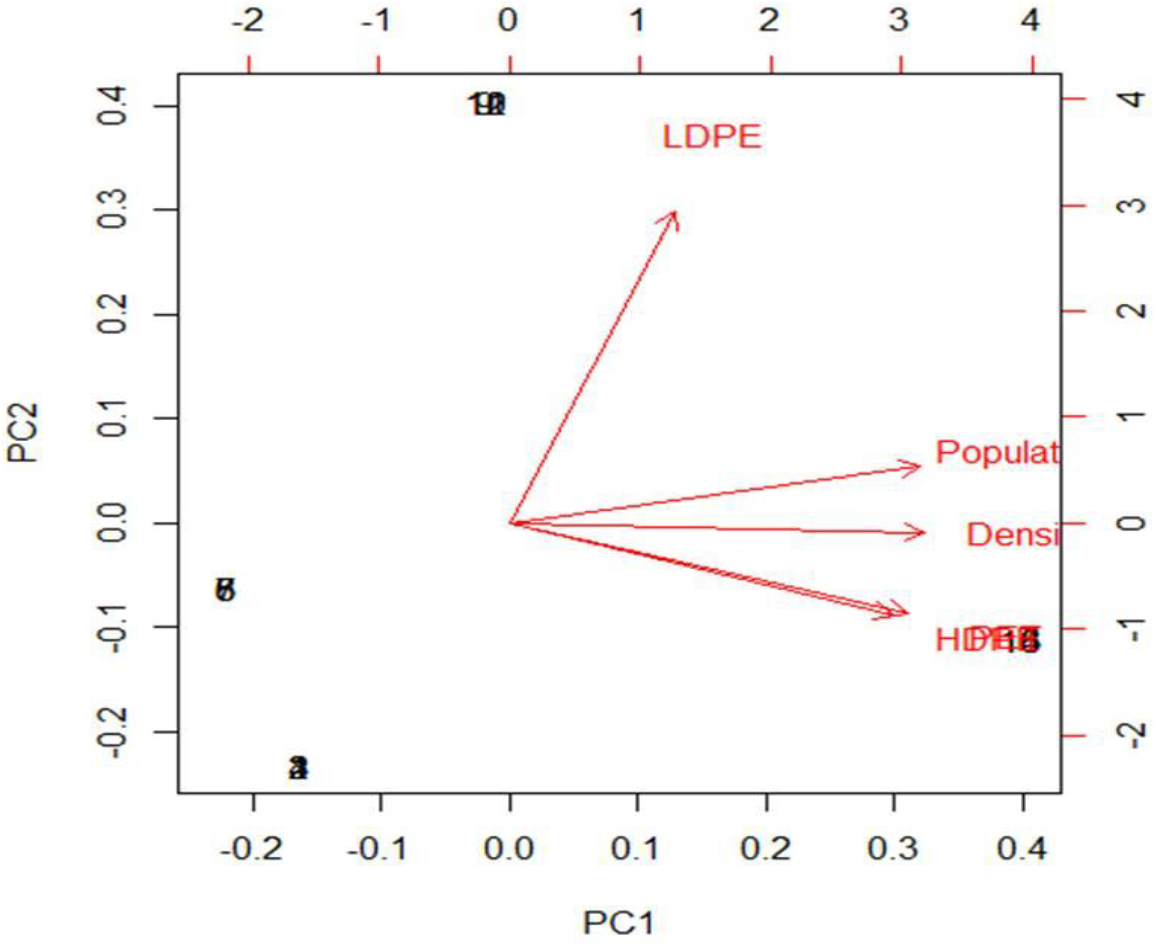
Principal Component Analysis of each PET, HDPE, and LDPE plastic litter density with population and population density covariates

## Discussion

High density of particular HDPE and LDPE litter in this study is related to the land use types and people activities (Aydın et al. 2016). Types of plastic litter released to the environment were influenced by wide ranges of land uses including the presences of industries, urbanized areas, and tourism-related activities (Yu et al. 2016). In almost all parts of Asia, polyethylene (PE) and polypropylene (PP) are the most common plastic litter. While in eastern and south-eastern Asia, Hong Kong, and Peninsular South Korea, expanded polystyrene (EPS) is more common (Fok & Cheung 2015, Kang et al. 2015, Kim et al. 2015). In particular Southeast Asia, Fauziah et al. (2015) have reported the abundance polystyrene (PS) and PE in the beaches of Peninsular Malaysia. LDPE is also common plastic litter in Asia. It has been reported from surface water of coastal in Hong Kong (Tsang et al. 2016) to lake (Sruthy & Ramasamy 2017).

The most common sources of plastic litters are including the tourism and fishery activities (Stolte et al. 2015), the release of wastewater from industries and urban run-off (Lima et al. 2015, Patters & Bratton 2016), and maritime transport activities (Gallagher et al. 2016, Veerasingam et al. 2016). However those possibilities can be excluded since the studied islands were small and do not have riverine system. There was a tourism activity whereas the magnitude of tourism is less significant. Regarding the industry effluent, the land uses in those islands were dominated by residential. The most possible of plastic litter discharge is due to direct littering since the study area is isolated. The similar situation is comparable to the presence of plastic litter in remote place. In isolated marine environment, Cincinelli et al. (2017) stated that the sources of plastic litter were related to the proximity to the local wastewater treatment plant, ship traffic, anthropogenic activities in the coastal area, and facilitated by transportation by means of ocean currents.

LDPE is the most common plastic litter and widely distributed across the surface water. In this study, LDPE is presence in the forms of plastic bags and this is the most common plastic materials and products used by the people every day. This wide distribution and uses of LDPE are explained by the population factor as the best model to estimate LDPE. It means that LDPE is used by the whole population in all islands in Thousand Island archipelago, while HDPE is only consumed by population living in island number 7. Among plastic types, LDPE demand is the highest accounted for 17.5% in comparison to HDPE, PET and other polymers. Besides high demand and lead to high discharges to the environment, high density of LDPE litter in the environment is also related to its persistence (Andrady 2015). This has made LDPE litter cannot be degraded at least in the short time and explains LDPE high density in this study. Since LDPE has no value and it is dumped and causing aesthetic, environmental, and public health issues (Kumi-Larbi et al 2016).

The second high plastic litter is the HDPE that commonly used as plastic wrap. High density of HDPE in surface water is related to the population density factors as estimated by the model. The contributions of population density on plastic litter is in accordance with other results from Chesapeake Bay (Yonkos et al 2014), Great Lakes (Baldwin et al. 2016), and urban wetlands (Townsend et al. 2019). Combined with the anthropogenic activities, population density will contribute more in increasing plastic quantities (Best 2019).

## Conclusion

Floating plastic litter density in the remote surface water of Southeast Asia is determined by several underlying factors. First, plastic litter spatial distribution is influenced by land use types. Second, the plastic litter density is a function of population demand and population density as a proxy of lack of available land that can be used to recycle the plastic litter and serves as landfill facilities. This becomes more imminent for the particular population that inhabits the isolated island with the only option is to discharge the litter directly to the nearby water body and marine ecosystem since all available land has been occupied and converted to settlement developments.

